# Effects of Slow Oscillatory Transcranial Alternating Current Stimulation on Motor Cortical Excitability Assessed by Transcranial Magnetic Stimulation

**DOI:** 10.1101/2021.05.13.444101

**Authors:** Asher Geffen, Nicholas Bland, Martin V Sale

## Abstract

Converging evidence suggests that transcranial alternating current stimulation (tACS) may entrain endogenous neural oscillations to match the frequency and phase of the exogenously applied current and this entrainment may outlast the stimulation (although only for a few oscillatory cycles following the cessation of stimulation). However, observing entrainment in the electroencephalograph (EEG) during stimulation is extremely difficult due to the presence of complex tACS artefacts. The present study assessed entrainment to slow oscillatory (SO) tACS by measuring motor cortical excitability across different oscillatory phases during (i.e., online) and outlasting (i.e., offline) stimulation. 30 healthy participants received 60 trials of intermittent SO tACS (0.75 Hz; 16s on / off interleaved) at an intensity of 2mA peak-to-peak. Motor cortical excitability was assessed using transcranial magnetic stimulation (TMS) of the hand region of the primary motor cortex (M1_HAND_) to induce motor evoked potentials (MEPs) in the contralateral thumb. MEPs were acquired at four time-points within each trial – early online, late online, early offline, and late offline – as well as at the start and end of the overall stimulation period (to probe longer-lasting aftereffects of tACS). A significant increase in MEP amplitude was observed from pre-to post-tACS (*P* = 0.013) and from the first to the last tACS block (*P* = 0.008). However, no phase-dependent modulation of excitability was observed. Therefore, although SO tACS had a facilitatory effect on motor cortical excitability that outlasted stimulation, there was no evidence supporting entrainment of endogenous oscillations as the underlying mechanism.

## 1 Introduction

Neural oscillations (i.e., cyclic fluctuations in neuronal excitability) are proposed to provide phase-dependent temporal regulation of neural information processing (Buzsáki 2006). In order to explore the functional relationships between neural oscillations and behaviour in normal brain function, rhythmic subtypes of transcranial electrical stimulation (tES) have been used to attempt to modulate endogenous neural oscillatory activity experimentally (Marshall & Binder 2013; Reato et al. 2013).

Converging evidence from both animal (Krause et al. 2019; Reato et al. 2013) and human studies (Helfrich et al. 2014a, b; Witkowski et al. 2016) suggests that during stimulation, tES may be able to entrain (i.e., synchronise) endogenous neural oscillations with respect to the frequency and phase of the exogenously applied current. Researchers have also been able to influence behavior by modulating the power of specific neural oscillations associated with behavioral outcomes (Antal & Paulus 2013; Ketz et al. 2018).

Unlike the well-documented immediate (online) effects of tES, there is less agreement regarding the magnitude and duration of post-stimulus (offline) effects (Veniero et al. 2015). These offline effects cannot be fully explained by a direct continuation of online entrainment (referred to as entrainment “echoes”), since these “echoes” only persist for a few oscillatory cycles following cessation of stimulation (Hanslmayr et al. 2014; Marshall et al. 2006; Thut et al. 2011). Therefore, longer-lasting offline effects (referred to as aftereffects) lasting up to 70 minutes are likely to reflect mechanisms other than entrainment *per se* (e.g. synaptic plasticity) (Kasten et al. 2016; Neuling et al. 2013; Veniero et al. 2015; Vossen et al. 2015).

The effects of tES on endogenous oscillatory activity have traditionally been quantified using electroencephalography (EEG; Jones et al. 2018; Ketz et al. 2018; Kirov et al. 2009; Marshall et al. 2006). However, observing entrainment in the EEG concurrently with tES is extremely difficult due to the presence of complex artefacts (Noury & Siegel 2017) and we have therefore used an alternative method to assess the effects of tES on cortical excitability. Single-pulse transcranial magnetic stimulation (TMS) is a form of non-invasive brain stimulation that can be used to indirectly probe the excitability of neocortical networks with high spatiotemporal precision of the order of millimetres and milliseconds (Hallett 2007). When applied to the hand area of the primary motor cortex (M1_HAND_), each TMS pulse induces a motor evoked potential (MEP) in the contralateral target muscle, the amplitude of which can then be measured using electromyography (EMG) (Barker et al. 1985). These MEP amplitudes provide an indirect measure of motor cortical excitability with good topographical specificity (Bergmann et al. 2012; Di Lazzaro et al. 2004; Hallett 2007; Ilmoniemi & Kicić 2010). By applying TMS pulses within a particular oscillatory phase of tES (referred to as phase-dependent stimulation), TMS can be used to assess entrainment of endogenous neural oscillations by tES (i.e., whether motor cortical excitability is modulated with respect to the phase of tES) (Raco et al. 2016; Schaworonkow et al. 2019; Zrenner et al. 2018). Importantly, the artefact issues of EEG are not present with TMS–EMG measures, thus allowing for an unambiguous investigation of tACS effects on motor cortical excitability.

Because we wanted to probe the phase-specific effects of tES, we chose to apply tES at a low frequency to allow MEP sampling across the different phases of stimulation. In this manner, the phase-cycle of low-frequency transcranial alternating current stimulation (tACS) could be conceptualised as representing alternating periods of classic ‘anodal’ and ‘cathodal’ transcranial direct current stimulation (tDCS), on which much earlier work has focused (Antal & Paulus 2013; Bland & Sale 2019; Liu et al. 2018; Reato et al. 2013). Therefore, in the present investigation, we chose to examine the online and offline effects of slow oscillatory (SO; 0.75Hz) tACS on motor cortical excitability using TMS.

Slow oscillations are typically prevalent during slow-wave sleep (SWS) and play an important role in sleep-dependent consolidation of motor learning (Marshall & Binder 2013). Despite the lack of endogenous SO activity during wakefulness, anodal SO tDCS during wakefulness has been shown to increase endogenous SO EEG power with relatively short-lasting offline effects (<1 minute) (Kirov et al. 2009), though the exact duration of these offline effects was not thoroughly assessed. However, it is important to note that these increases in SO power were more restricted topographically to the prefrontal cortex (the predominant source of endogenous slow oscillations during sleep) and were less pronounced than those observed following SO tDCS applied *during* SWS (Marshall et al. 2006). Furthermore, due to the previously mentioned complexity of tES artefacts in the EEG, the authors could not determine whether these localised increases in EEG SO power were in fact due to the entrainment of slow oscillations by SO tDCS. Therefore, it remains unclear whether slow oscillations can be reliably entrained in the awake brain at intensities typical of tES (i.e., 1–2 mA).

Anodal SO tDCS during wakefulness has also been shown to induce lasting increases in motor cortical excitability that persist beyond stimulation (Bergmann et al. 2009; Groppa et al. 2010). However, due to the anodal component (i.e., positive current offset) of this stimulation—which in itself can cause an increase in cortical excitability (Bergmann et al. 2009; Nitsche & Paulus 2000; Nitsche et al. 2007)—it cannot be concluded that these effects are a direct result of the influence of the applied slow oscillations. tACS has a significant technical advantage over tDCS in this regard, since it has no DC offset (i.e., an average current of 0 mA). Despite this, the effects of SO tACS on motor cortical excitability have not been thoroughly examined in previous literature. Antal et al. (2008) found no significant changes in motor cortical excitability following SO (1 Hz) tACS; however, their stimulation protocol was suboptimal for inducing changes in endogenous oscillatory activity due to the low stimulation intensity (0.4 mA; Huang et al. 2008; Reato et al. 2010) and constant rather than intermittent application of tACS (Jones et al. 2018; Ketz et al. 2018).

The aims of this study were (1) to investigate the online effects of SO tACS applied intermittently at high intensity on motor cortical excitability; (2) to determine if tACS-induced changes in motor cortical excitability persist beyond each trial of stimulation (i.e., entrainment echoes) as well as beyond the total stimulation period (i.e., offline aftereffects).

It was hypothesised that SO tACS will induce SO-like sinusoidal changes in motor cortical excitability that correspond with the tACS phase, with high MEP amplitudes at oscillatory peaks and low amplitudes at oscillatory troughs, supporting online entrainment. Secondly, that sinusoidal changes in motor cortical excitability will persist for a few oscillatory cycles immediately following each trial of stimulation, thus demonstrating entrainment echoes. Thirdly, that motor cortical excitability will increase over the total duration of stimulation (although this relationship may not necessarily be linear), and this increase will be sustained beyond the total stimulation period (i.e., offline aftereffects).

## 2 Materials and Methods

### 2.1 Subjects

Forty-one healthy, right-handed participants (17 male, aged 24 ± 4 years) were recruited by advertisement, although 11 participants were excluded from the final analysis (see “MEP screening” below) leaving a sample size of 30 participants. All participants completed a safety screening questionnaire (Keel et al. 2001) and provided a written statement of informed consent prior to commencing the experiment. Approval was granted by the University of Queensland Human Research Ethics Committee.

### 2.2 Quantification of Motor Cortical Excitability Using TMS

Motor cortical excitability was assessed by measuring TMS-induced MEP amplitudes that were recorded from the target muscle using surface EMG. The target site for the TMS was the left M1_HAND_ region, specifically the region associated with the *abductor pollicis brevis* (APB), a large thumb muscle.

### 2.3 Experimental Setup

#### 2.3.1 EMG

Participants were seated comfortably in a chair and their right forearm placed on a foam mat with their forearm supinated. EMG activity of the APB muscle was recorded using disposable surface electrodes (H124SG 30mm x 24mm). Positive and negative electrodes were placed over the APB muscle belly and at the first metacarpophalangeal joint respectively and a reference electrode was placed on the anterior surface of the wrist.

#### 2.3.2 TMS

TMS pulses were applied to the left M1_HAND_ region using a Magstim Double 70mm Remote Control Coil charged by a Magstim 200^2^ stimulator (Magstim, UK). The individual location of the left M1_HAND_ region as well as the TMS intensity (resting motor threshold – RMT) were determined for each participant using manual TMS “hot-spotting” (Rossini et al. 1994). This involves systematically adjusting the position of the TMS coil on the participant’s head whilst also adjusting the stimulation intensity until MEPs are consistently induced (i.e., in at least five out of ten successive trials) with amplitudes above a desired threshold (0.5 mV). The location of the left M1_HAND_ region was then marked on the participant’s scalp using an erasable marker.

#### 2.3.3 tACS

tACS was applied using a NeuroConn DC Stimulator Plus. The 42×45mm tACS pad electrodes were applied to the scalp over the left motor cortex and the contralateral supraorbital region. Before attaching the electrodes, the scalp was rubbed with ethanol (70%) and Ten20 conductive paste was applied to the electrodes to minimise resistance between the electrodes and scalp. The electrode targeting the left motor cortex was not placed on the marked hotspot itself, but rather ∼2 centimetres posterolateral to the hotspot. This slight increase in inter-electrode distance is thought to reduce current shunting through the scalp and cerebrospinal fluid, thus, maximising current density at the target site and increasing the effectiveness of the tACS (Faria et al. 2011).

#### 2.3.4 Recording tACS Output

The tACS output was recorded using disposable surface electrodes (H124SG 30mm x 24mm). These electrodes were placed over the tACS pad targeting the supraorbital region and on the left side of the forehead, and referenced to the nose tip. The tACS artefact was used to synchronise stimulation with the computer used for MEP acquisition (i.e., such that each probe by TMS was timed with respect to the phase of tACS).

#### 2.3.5 Data Collection

All surface electrode measurements (i.e., APB EMG and tACS output electrodes) were acquired (1KHz sampling rate; 20-1000Hz band pass filtering) via an electrode adaptor (Model CED1902-1 ½), before being amplified by a CED1902, and finally recorded by a CED1401 MICRO3 (Cambridge Electronic Designs, Cambridge, UK). TMS triggers were directly recorded by the CED1401 MICRO3. All data were then transferred from the CED1401 MICRO3 to a PC and saved via Signal (Ver. 6.04) software (Cambridge Electronic Designs, Cambridge, UK), before being exported to MATLAB (Ver. R2019a) for analysis.

## 2.4 Experimental Procedure

### 2.4.1 tACS Paradigm

Participants received 60 trials of tACS, with each trial consisting of 16 seconds (12 cycles at 0.75Hz) of tACS at an intensity of 2mA (“Online”), followed by 16 seconds of no tACS (“Offline”), for a total of 16 minutes of tACS and 16 minutes of no tACS (Figure 1A). The entire stimulation period was divided into 3 blocks (∼10 minutes comprising 20 trials each), with 5-minute rest periods (no tACS or TMS delivered) between blocks.

**Figure 1.**
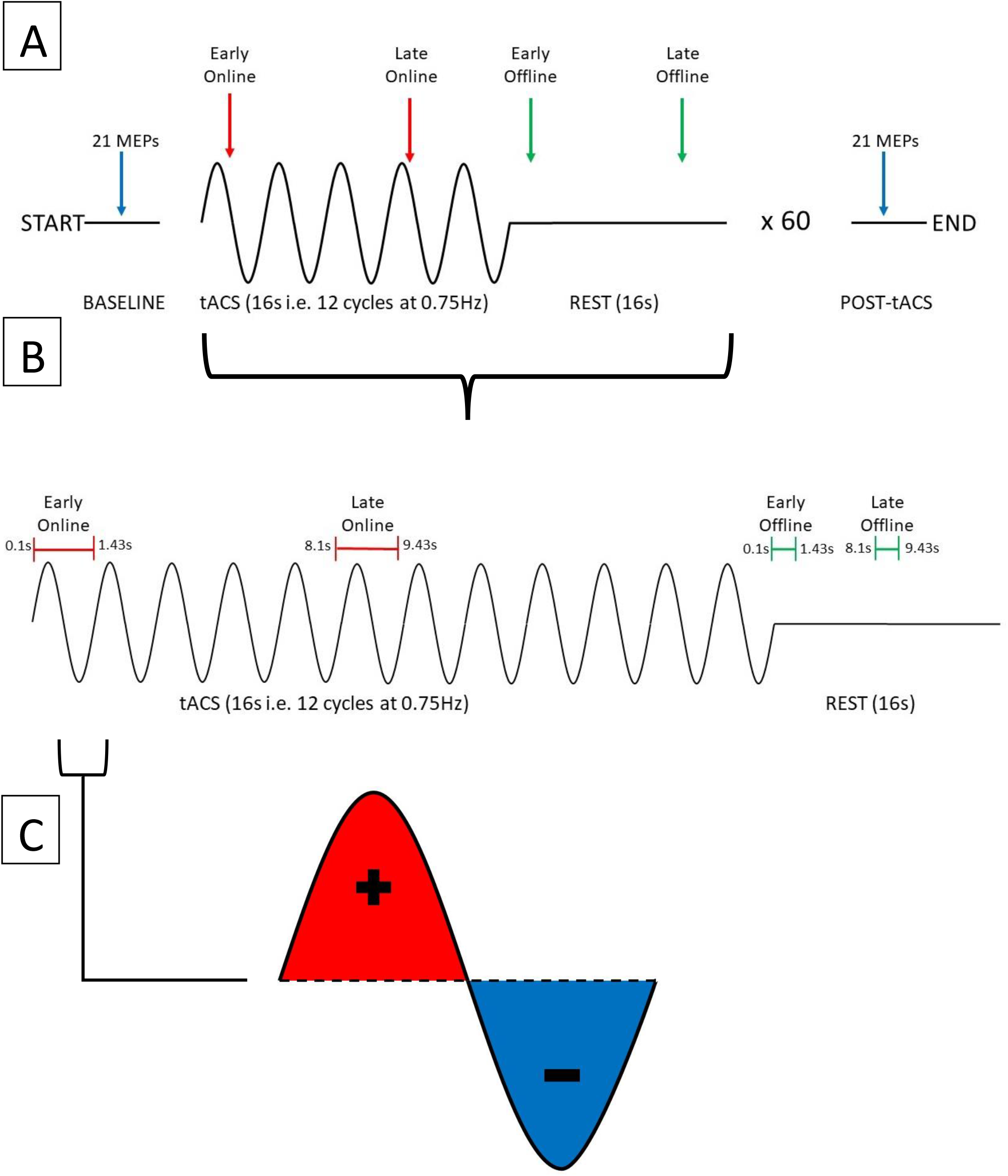
Summary of experimental procedure for probing changes in motor cortical excitability induced by SO tACS. A) The experimental session consisted of a 16-second tACS period (represented as sine waves) followed by a 16-second rest period (represented as flat lines), repeated 60 times for a total of ∼32 minutes. To probe how tACS-induced changes in motor cortical excitability evolve over time, 21 TMS-induced MEPs were acquired at the start and the end of the experimental session (blue arrows), 2 MEPs were acquired at each tACS period (red arrows), and 2 MEPs were acquired at each rest period (green arrows). B) Within each tACS trial (i.e., tACS + rest period), MEPs were acquired at 4 distinct time-points (1 MEP per time-point): early online, late online, early offline, and late offline. C) tACS alternates polarity between the anode and cathode to produce a sinusoidal current with both positive (red) and negative (blue) peaks.

### 2.4.2 TMS Paradigm

To examine the online and offline effects of SO tACS on motor cortical excitability, MEPs were acquired at 4 time-points within each trial (1 MEP per time-point): early online (0.1-1.43s after tACS starts), late online (8.1-9.43s after tACS starts), early offline (0.1-1.43s after tACS ends), and late offline (8.1-9.43s after tACS ends) (Figure 1B). Therefore, 60 MEPs were acquired for each time-point (i.e., once each per trial).

To examine if the effects of SO tACS on motor cortical excitability are specific to the tACS phase (both online and offline), sufficient MEPs (i.e., >20) need to be acquired across the different phases of the tACS (Goldsworthy et al. 2016). This was achieved by implementing a “jitter” (i.e., a randomised time delay) to the delivery of TMS so that the delivery was not locked to a specific phase of the tACS, and thus, TMS pulses were approximately uniformly delivered across the different phases across the entire stimulation block (Figure 1C).

To examine the cumulative effects of the entire tACS paradigm on motor cortical excitability, 21 TMS-induced MEPs were acquired both at baseline and at the end of the entire period of tACS delivery (Figure 1A). TMS was delivered every 0.2Hz.

## 2.5 Statistical Analysis

### 2.5.1 Data Transformation

The first MEP for each data set (as well as the first MEP after each of the rest periods) was always excluded, since initial MEP amplitudes may be larger (Brasil-Neto et al. 1994) and more variable (Schmidt et al. 2009) than subsequent MEPs, which can impact the reliability of TMS measures of cortical excitability. Further, individual MEPs were excluded if voluntary pre-MEP EMG activity was detected in the 500ms prior to TMS delivery (2.72% of MEPs excluded). Finally, participants with mean pre-tACS amplitudes less than 0.5mV or greater than 1.5mV were excluded from the final analysis (11 participants excluded). This is because excessively small or large pre-tACS MEPs may have introduced floor and ceiling effects, respectively (Cuypers et al. 2014).

The TMS triggers were then automatically categorised into their respective time-points (i.e., early online, late online, early offline, and late offline – see Figure 1B) and the “late online” triggers were used to calculate the tACS phase, since these triggers are the only ones where tACS was present both before and after TMS was applied, thus, providing the most reliable estimate of tACS phase. If tACS-induced phase entrainment persists beyond stimulation, we would expect the tACS phase to continue into the offline period. To assess this, the computed phase for the late online triggers was extrapolated (both forward and backward) and its values computed at each of the other time-points (Figure 2).

**Figure 2.**
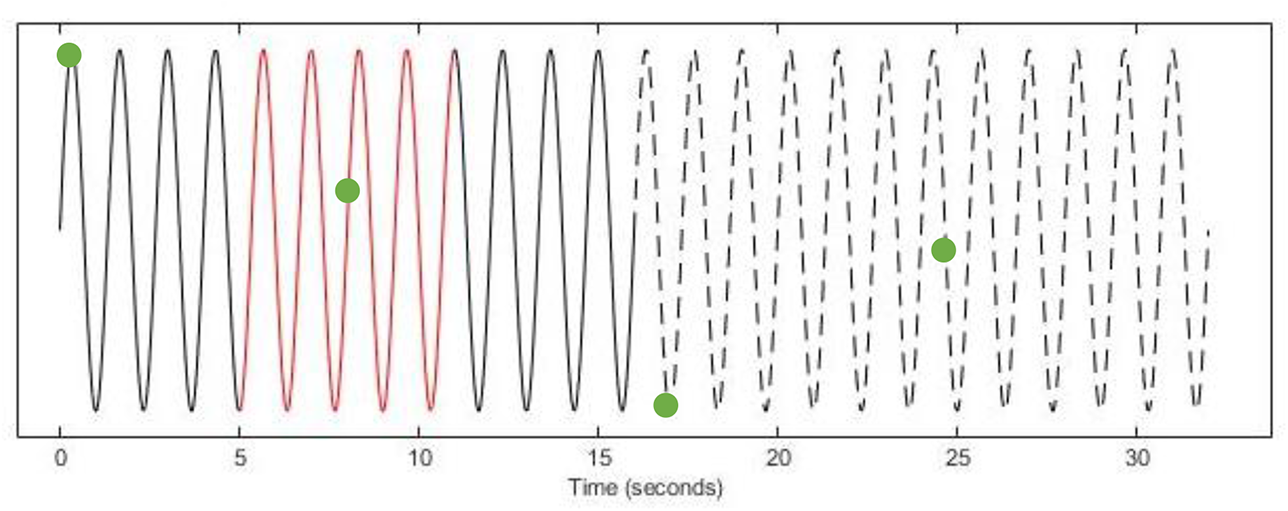
Determining tACS Phase at TMS Triggers (EXAMPLE ONLY). Using the tACS-output recording (Solid Line), a 6-second window (Red) of the instantaneous phase (centred on each late online trigger) was computed. The computed phase was then extrapolated both forward into the offline period (Dashed Line) and backward to the early online triggers (Solid Line) and the phase was computed at each of the other time-points (Green Dots).

### 2.5.2 Data Analysis

To determine if the effects of SO tACS on motor cortical excitability are specific to the tACS phase a permutation analysis was performed (Zoefel et al. 2019). This analysis was performed separately for each of the four time-points (∼60 MEPs per time-point per participant) as well as for all online and offline MEPs (∼120 MEPs online/offline per participant).

For the permutation analysis, an ideal sinusoidal model (0.75 Hz) was fitted to each participant’s observed MEP amplitudes for each of the four time-points based on their phase (Bland & Sale, 2019), and the amplitudes of these models were summed. The MEP amplitudes were then shuffled with respect to their phases. Next, ideal sinusoidal models were fitted to the shuffled data, and the amplitudes of these shuffled models were summed. This process was repeated for a total of 1000 permutations per participant. The true and shuffled summed amplitudes were then compared. In this analysis, the *P*-value is the proportion of shuffled summed amplitudes exceeding the true sum of amplitudes, remembering that under the null hypothesis (which assumes that the observed effects are not phase-specific) the amplitude of these sinusoidal models should be small (i.e., closer to zero). Because the permutation procedure disrupts any phasic effects that may be present, the shuffled MEPs act as a negative control for the true MEPs, and thus, the permutation analysis does not require a sham stimulation condition as a negative control.

To determine if there was a significant difference in mean MEP amplitudes between the pre- and post-tACS measurements, a paired sample *t*-test was performed with *P*-value < 0.05 considered significant. To examine how changes in MEP amplitudes evolve throughout the tACS period, additional paired sample *t*-tests were performed comparing each of the 3 tACS “blocks” against each other, again with *P*-value < 0.05 considered significant.

## 3 Results

### 3.1 Phase-Specificity of tACS Effects

The phase-specificity of acute changes in motor cortical excitability induced by SO tACS was assessed by a permutation analysis. Ideal sinusoidal models were fitted to each participant’s observed MEP amplitudes for each TMS time-point (∼60 MEPs per time-point per participant) based on their phase and the amplitudes of these models were summed. An example of one of these fitted sinusoidal models is shown in Figure 3. The true sum of amplitudes was then compared against the summed amplitudes of 1000 permutations of the MEP amplitudes, with *P*-values representing the proportion of shuffled summed amplitudes exceeding the true sum of amplitudes. The permutation analysis did not reveal significant phase-specific modulation of motor cortical excitability at any of the four TMS time-points (*P* = 0.86, 0.81, 0.21, and 0.70 for early online, late online, early offline, and late offline respectively). Combining the early and late online/offline MEPs (∼120 MEPs online/offline per participant) also failed to reveal any significant phase-specific modulation of motor cortical excitability online or offline to tACS (*P* = 0.89 and 0.90 respectively).

**Figure 3.**
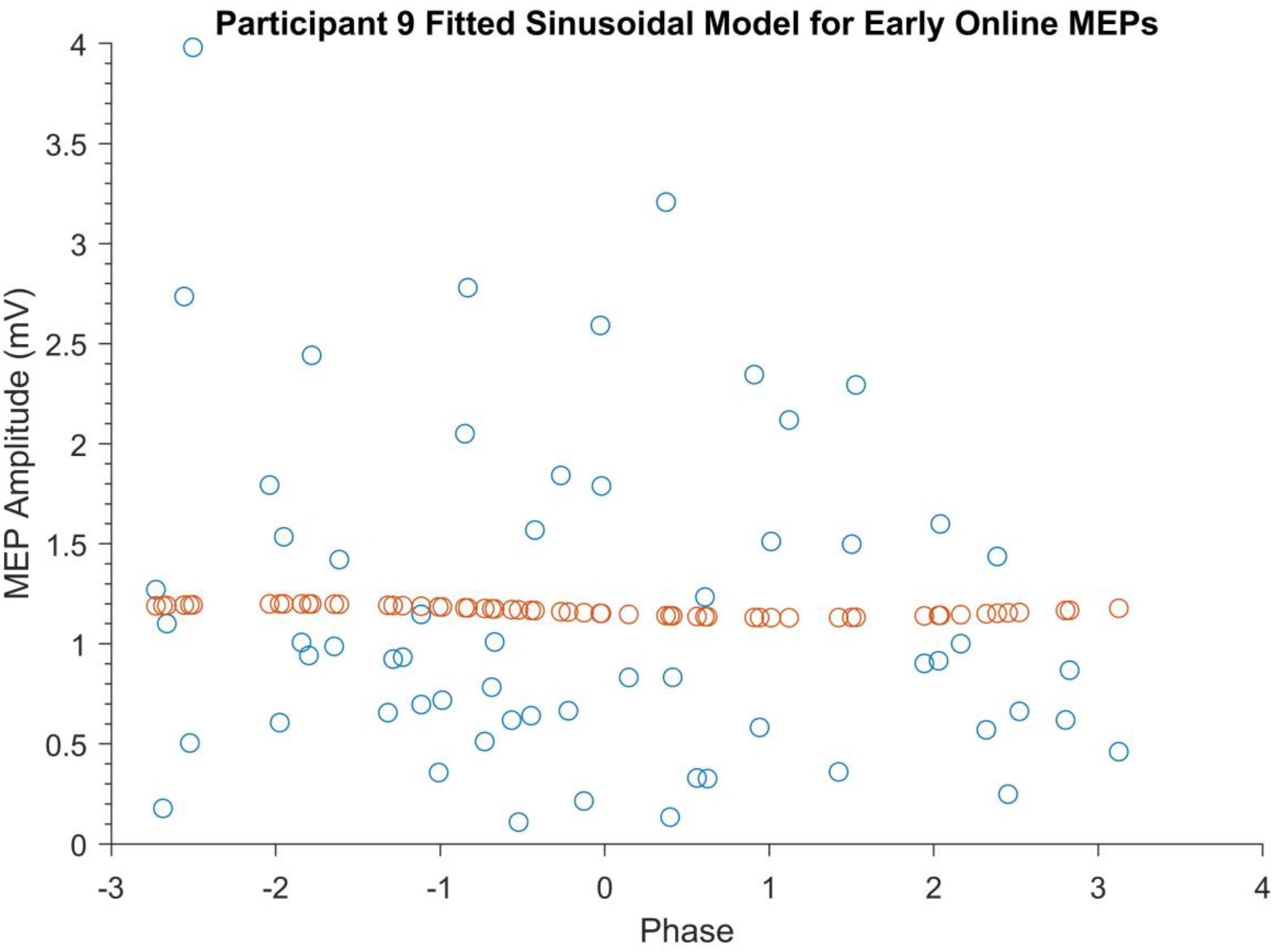
Example of a Participant’s Fitted Sinusoidal Model for Early Online MEPs. Blue dots represent Participant 9’s early online MEPs sorted according to tACS phase. Orange dots represent the fitted sinusoidal model for these MEPs.

### 3.2 Cumulative Effects of SO tACS on Motor Cortical Excitability

The cumulative effects of the tACS paradigm on motor cortical excitability was assessed by comparing mean MEP amplitudes pre- and post-tACS. As shown in Figure 4, MEP amplitudes were found to be significantly greater post-tACS (mean = 1.24mV ± 0.83) compared to pre-tACS (mean = 0.88mV ± 0.25) (*P* = 0.013).

**Figure 4.**
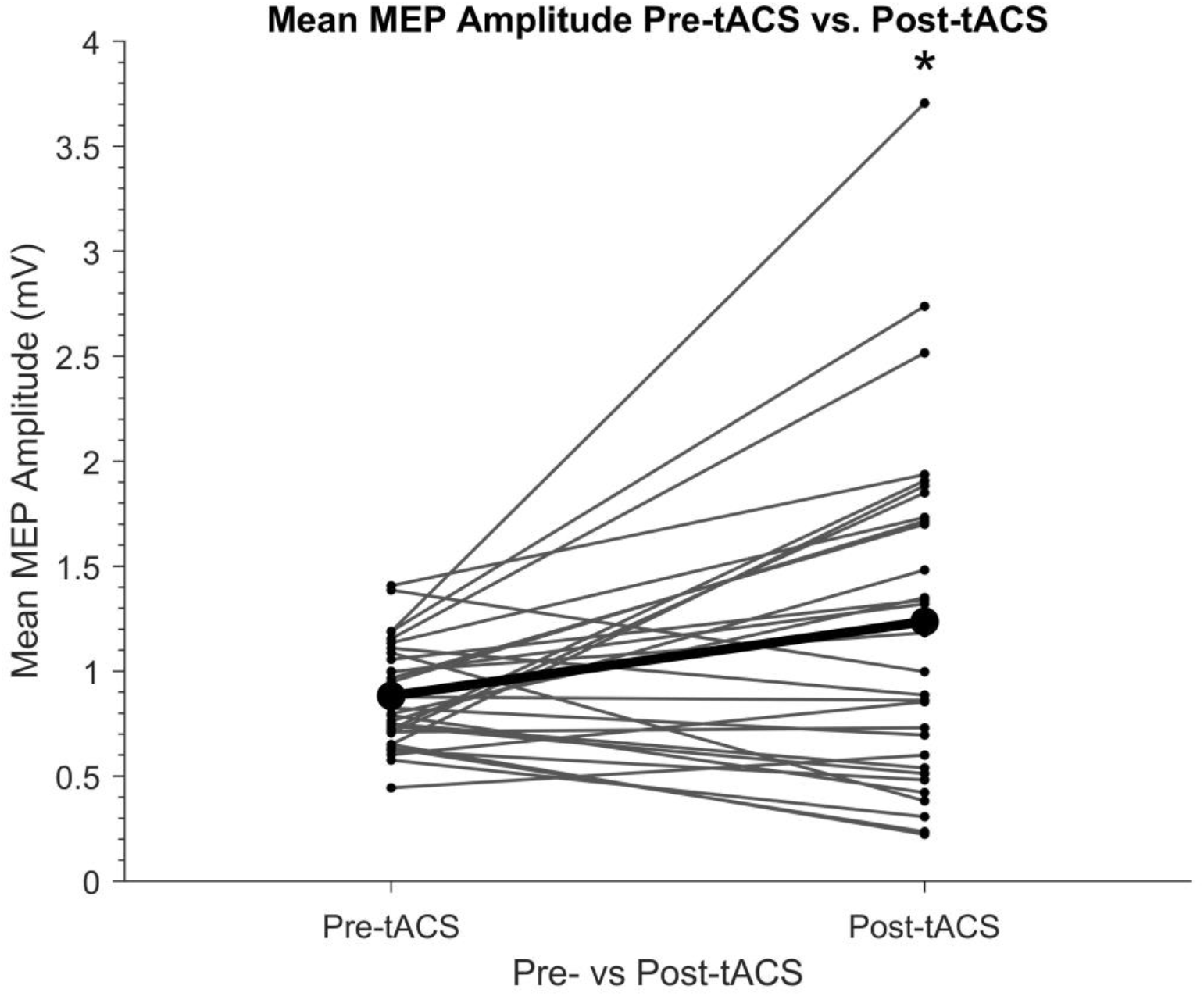
Individual Mean Amplitudes of TMS-induced MEPs Before and After SO tACS. Points represent each participant’s mean MEP amplitude from 20 TMS-induced MEPs (per participant) acquired before (pre-tACS) and after (post-tACS) receiving 60 “trials” (∼32 minutes) of SO tACS, with each trial consisting of 16 seconds of tACS (12 cycles at 0.75Hz) followed by 16 seconds of rest. Individual differences in mean MEP amplitude are represented by the solid lines. The global mean MEP amplitude for each group is represented by a bold point, with the global mean difference from pre-to post-tACS represented by a bold line connecting the two bold points. A significant increase in MEP amplitude from pre-tACS to post-tACS was reported (paired sample t-test P = 0.013; *). n = 30.

Because there was a significant increase in mean MEP amplitude from pre-to post-tACS, the question arose of whether this overall change in MEP amplitude occurred gradually over time within the stimulation period. We therefore compared mean MEP amplitudes within each of the three tACS blocks. As shown in Figure 5, there was a significant increase in MEP amplitudes between the 1^st^ (mean = 1.19mV ± 0.72) and 3^rd^ (mean = 1.46mV ± 0.94) blocks (*P* = 0.008), whereas there were no significant differences between the 1^st^ and 2^nd^ (mean = 1.33mV ± 0.79) blocks or between the 2^nd^ and 3^rd^ blocks (*P* = 0.12 and 0.33 respectively). This suggests a gradual build-up of cortical excitability from tACS.

**Figure 5.**
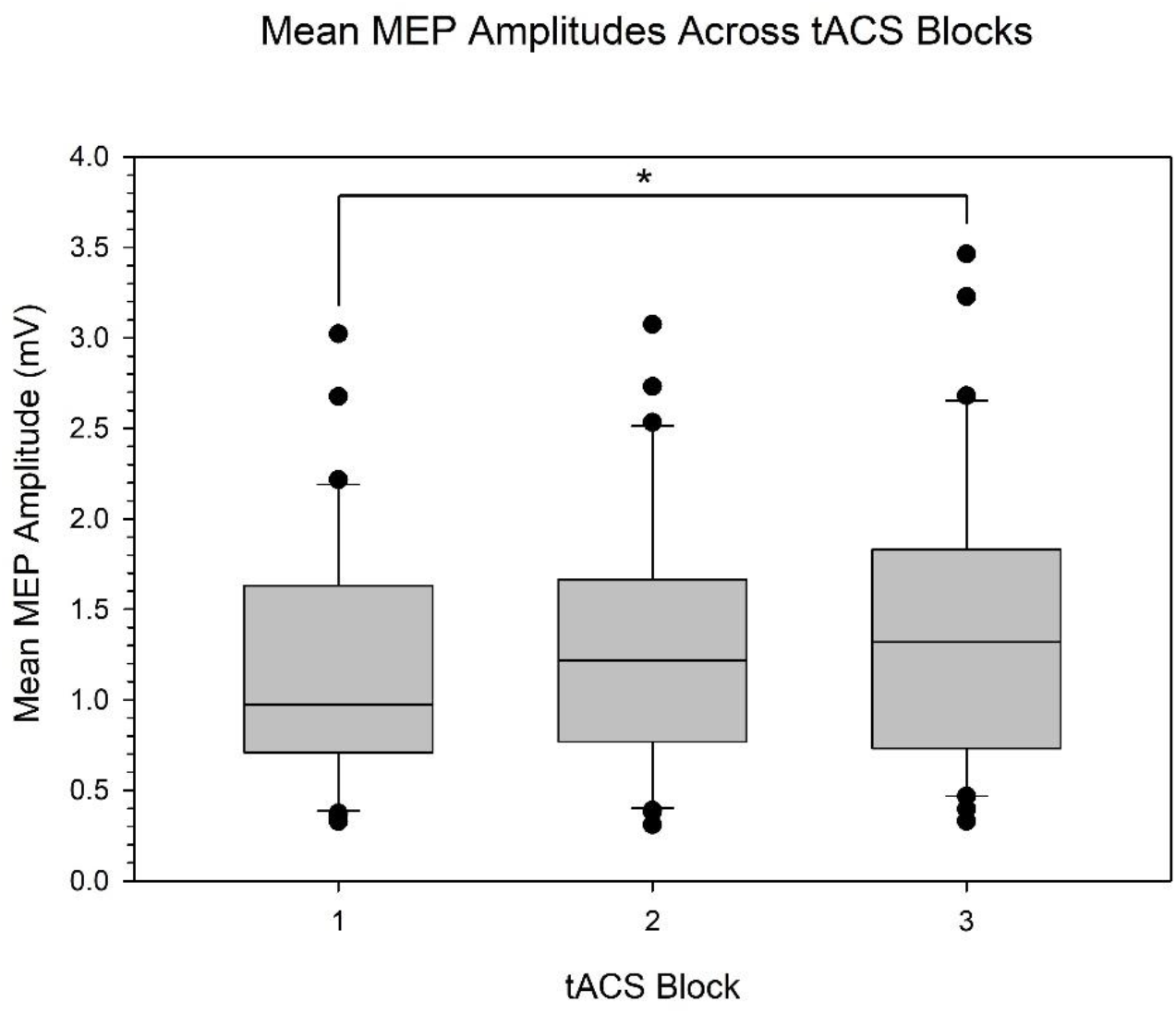
Boxplots of TMS-induced MEP Amplitudes Within Each Block of the tACS Period. Boxes represent the IQR of TMS-induced MEP amplitudes from ∼80 MEPs (per participant) acquired during 3 “blocks” of a ∼32-minute tACS period. Each block contains ∼20 “trials” (∼10 minutes) of SO tACS, with each trial consisting of 16 seconds of tACS (12 cycles at 0.75Hz) followed by 16 seconds of rest. The horizontal black line within each box indicates the median and the whiskers below and above each box indicate the 10^th^ and 90^th^ percentiles respectively. Outliers beyond the 10^th^ and 90^th^ percentiles are denoted by black circles. No significant differences in MEP amplitude were observed between the 1^st^ and 2^nd^ blocks (paired sample t-test P = 0.12) or the 2^nd^ and 3^rd^ blocks (P = 0.33). However, a significant difference in MEP amplitude between the 1^st^ and 3^rd^ blocks was observed (P = 0.008; *). n = 30.

It is worth mentioning that widening the exclusion criteria for participants based on their pre-tACS mean MEP amplitudes (0.5–1.5 mV to 0.4–2 mV) did not affect the significance of the cumulative or phase-specific effects of SO tACS in the present study, despite increasing the sample size from 30 to 36 participants.

## 4 Discussion

Although there has been a plethora of studies in the last decade reporting behavioural, perceptual, and electrophysiological effects induced by tACS (Antal & Paulus 2013, Bland & Sale 2019, Liu et al. 2018), the mechanisms underlying these effects remain controversial. In the present study, we aimed to probe SO tACS-induced entrainment of endogenous slow oscillations both online and offline by assessing motor cortical excitability across different oscillatory phases using TMS-induced MEP amplitudes. We also assessed the cumulative effects of SO tACS on motor cortical excitability by comparing mean excitability pre- and post-stimulation as well as comparing mean excitability across stimulation blocks.

Regarding the cumulative effects of SO tACS, we present the first evidence of enhanced motor cortical excitability induced by SO tACS in the awake brain, with a significant increase in TMS-induced MEP amplitudes from pre-to post-tACS as well as from the 1^st^ to the 3^rd^ tACS block. Excitability increases induced by anodal SO tDCS have been reported previously (Bergmann et al. 2009, Groppa et al. 2010). However, compared to the SO tDCS study by Bergmann et al. (2009), the present study using SO tACS demonstrated a greater MEP amplitude increase (40.91% vs. 22%) despite shorter stimulation periods (16s vs. 30s), shorter total duration of stimulation (16min vs 17.5min), longer rest periods between trials (16s vs. 5s) and two additional 5-minute rest periods. Critically, due to the lack of an anodal component for tACS, this excitatory effect cannot be attributed to a general depolarisation of cortical motor neurons and is thus driven by some other factor.

It is theoretically possible that the cumulative increase in motor cortical excitability was associated with an entrainment of endogenous slow oscillations (Jones et al. 2018; Ketz et al. 2018; Kirov et al. 2009; Marshall et al. 2006). However, the acute effects of stimulation did not appear to be dependent on the tACS phase, with the permutation analysis providing no significant evidence for phase-specific modulation of motor cortical excitability at any of the four TMS time-points (i.e., early/late online/offline) or for the combined online/offline MEPs.

The most likely explanation for the lack of an entrainment effect in the present results is that endogenous slow oscillations are not prevalent enough in the wake brain to be effectively entrained by SO tACS. This conclusion is in line with previous SO tDCS/tACS studies (Bergmann et al. 2009; Groppa et al. 2010; Jones et al. 2018; Ketz et al. 2018; Kirov et al. 2009; Marshall et al. 2006) as well as tACS studies using different stimulation frequencies (Ali et al. 2013; Antal et al. 2008; Kanai et al. 2008) that found stimulation to be most effective when the frequency of the exogenously applied oscillations closely matches the frequency of the predominant endogenous oscillations. These findings suggest that network resonance is a key underlying mechanism by which tACS modulates large-scale cortical network activity. These resonance dynamics are characterised by a phenomenon called an “Arnold Tongue”, where the current intensity required to induce a particular oscillation increases the more the frequency of that oscillation deviates from the resonant (eigen) frequency of the network (Ali et al. 2013; Liu et al. 2018; Thut et al. 2017).

In future experiments, tACS will instead be applied at a frequency that is naturally present in the motor cortex during wakefulness, such as mu (*μ)* rhythms (∼9-11Hz) (Antal et al. 2008; Feurra et al. 2019; Gundlach et al. 2017; Madsen et al. 2019; Thies et al. 2018; Wach et al. 2013). This will also allow us to determine if the excitatory effect observed in the present experiment is specific to SO tACS or if similar effects are observed for other stimulation frequencies.

Alternatively, because the present experiment did not include a negative control stimulation condition for SO tACS (e.g., sham stimulation), it is theoretically possible that the observed increase in motor cortical excitability is simply a time-dependent effect and not a true tACS effect. However, this is highly unlikely given that a recent meta-analysis by Dissanayaka et al. (2018) did not report any significant effects of sham tES on cortical excitability compared to baseline, even up to 90 minutes following sham tES (Chaieb et al. 2011; Moliadze et al. 2010, 2012).

Although the underlying cause of the observed increase in MEP amplitude cannot be concluded from the present results, the lack of phase-specific entrainment suggests that this excitatory effect may instead be driven by plasticity-related mechanisms, such as spike-timing dependent plasticity (STDP) (Veniero et al. 2015; Vossen et al. 2015). In the STDP model, even a slight mismatch between the stimulation frequency and an individual’s spontaneous peak frequency could influence the direction of any induced changes, which may explain the heterogeneity of tACS after-effects across studies (Veniero et al. 2015). Tests of this model should therefore tailor stimulation frequency to each participant’s individual peak frequency rather than use a standard frequency such as was used in the present study.

It is important to note that entrainment and plastic-like effects induced by tACS are not mutually exclusive (Vosskuhl et al. 2018). In fact, Helfrich et al. (2014a,b) found the magnitude of induced aftereffects to be positively correlated with the magnitude of online entrainment and also demonstrated that online effects occurred within a narrow frequency peak whilst offline effects occurred across a broader band around this peak; thus, eliminating the possibility that the online and offline effects reflect the same phenomenon at different points in time.

Elucidation of the mechanisms underlying the online and offline effects of tACS will enable its therapeutic applications. For example, the ability to induce lasting plastic changes in the motor cortex using tACS may improve the effectiveness of existing rehabilitation for neurological conditions where motor function is compromised.

## 5 Conclusion

In summary, the significant increase in TMS-induced MEP amplitudes from pre-to post-SO tACS as well as from the 1^st^ to the 3^rd^ SO tACS block suggests that, similar to previously reported excitability increases induced by anodal SO tDCS, SO tACS had a facilitatory effect on motor cortical excitability that outlasted the stimulation period. Importantly, the present findings suggest that these motor cortical excitability increases are not simply due to anodal stimulation. However, given the acute effects of SO tACS were independent of phase, this study does not support entrainment of endogenous slow oscillations as an underlying mechanism for this excitatory effect.

## Acknowledgements

We would like to thank Dr. Kylie Tucker for her role as co-supervisor during Asher Geffen’s honours research program. We would also like tothank Professors Laurie and Gina Geffen for their help in proofreading this manuscript.

## Abbreviations

tES: Transcranial Electrical Stimulation
tACS: Transcranial Alternating Current Stimulation
tDCS: Transcranial Direct Current Stimulation
SO: Slow Oscillatory
SWS: Slow-Wave Sleep
TMS: Transcranial Magnetic Stimulation
EEG: Electroencephalography
EMG: Electromyography
MEP: Motor Evoked Potential
M1_HAND_: Hand Area of the Primary Motor Cortex
APB: *abductor pollicis brevis*
STDP: Spike-timing dependent plasticity

## Graphical Abstract

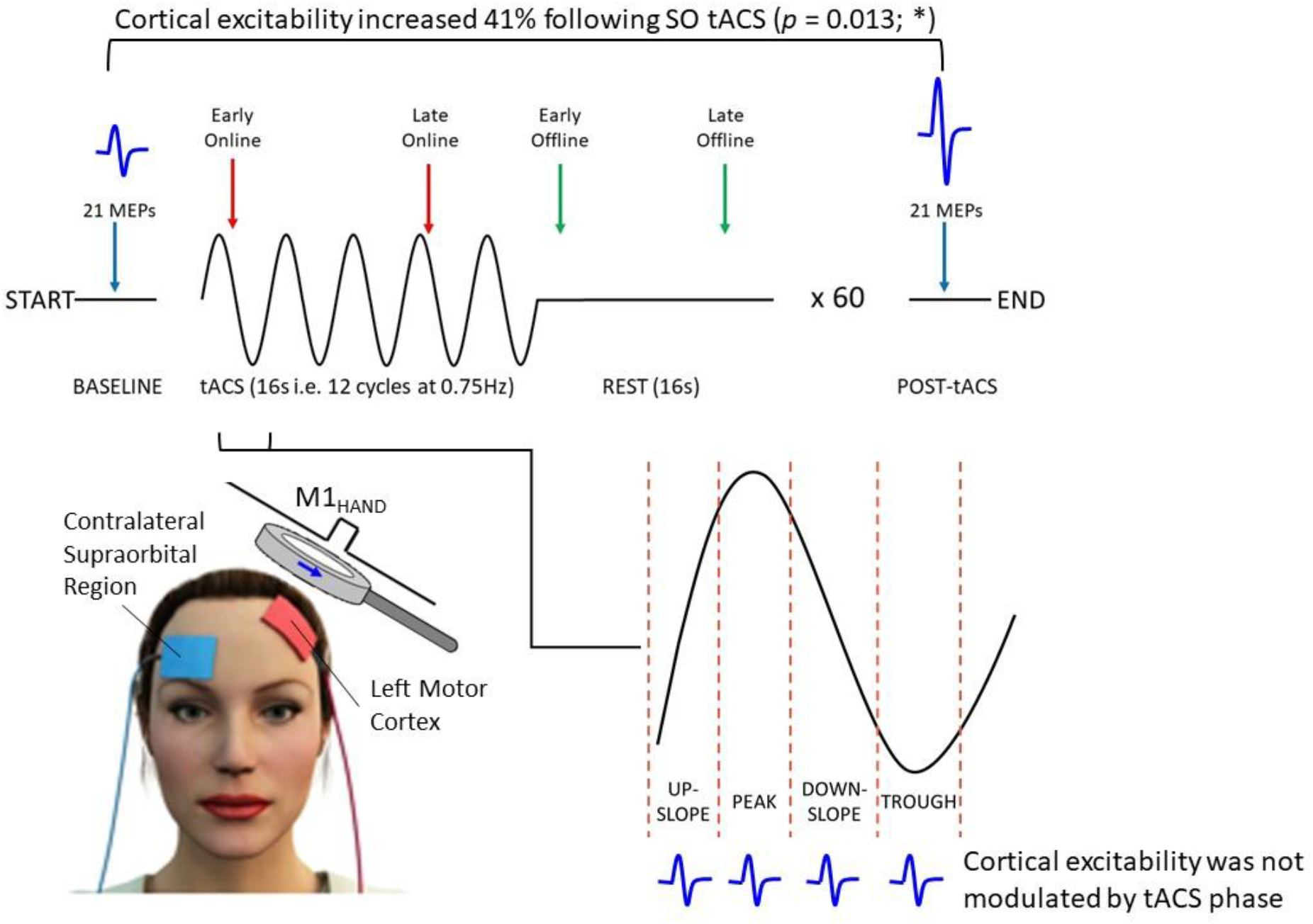

30 healthy participants received 60 trials of intermittent SO (0.75 Hz) tACS (1 trial = 16s on + 16s off) at an intensity of 2mA. Motor cortical excitability was assessed using TMS-induced MEPs (blue waveforms) acquired across different oscillatory phases during (i.e., online; red arrows) and outlasting (i.e., offline; green arrows) tACS, as well as at the start and end of the stimulation session (blue arrows). Mean MEP amplitude increased by ∼41% from pre-to post-tACS (*P* = 0.013); however, MEP amplitudes were not modulated with respect to the tACS phase.

## Notes

<u>Funding:</u> This work was supported by the US Office of Naval Research Global [grant number N62909-17-1-2139] awarded to Martin V Sale. The funding body had no involvement in the study design; the collection, analysis, and interpretation of data; the writing of the report; or the decision to submit the article for publication.

<u>Conflicts of interest:</u> The authors have no relevant financial or non-financial interests to disclose.

### Competing Interest Statement

The authors have declared no competing interest.

